# Evaluation of *IGF1* and *MKI67* relative abundance of mRNA transcript in relation to histopathological features of equine endometrosis

**DOI:** 10.64898/2026.02.23.707461

**Authors:** Łukasz Zdrojkowski, Anna Niwińska, Ewa Kautz, Dawid Tobolski, Magdalena Fajkowska, Małgorzata Rzepkowska, Tomasz Jasiński, Małgorzata Domino, Bartosz Pawliński

**Author notes:** Corresponding author (ŁZ).

## Abstract

Equine endometrosis is a major cause of subfertility in mares characterized by fibrotic remodeling of the endometrium. Present study evaluated endometrial relative abundance of mRNA transcript of *IGF1, MKI67, TGFB1*, and *ACTA2* in relation to endometrosis severity and defined histopathological features.

Forty-seven endometrial samples were graded according to the modified Kenney and Doig (KD) categories. Relative abundance of mRNA transcript was quantified by RT-qPCR and histopathology was extended using a standardized feature-based assessment.

*TGFB1* relative abundance of mRNA transcript was higher in category I+ than in category I and in samples with glandular basal lamina disruption. *MKI67* relative abundance of mRNA transcript was lower in samples with luminal epithelial erosion. *IGF1* relative abundance of mRNA transcript correlated negatively with KD category, glandular degeneration, overall inflammatory infiltration, lymphocytic infiltration, and neutrophilic infiltration. *MKI67* correlated positively with *ESR1*.

These findings indicate that early endometrosis-compatible lesions are associated with increased *TGFB1* transcription and that epithelial damage is accompanied by reduced *MKI67* abundance of mRNA. The inverse associations between *IGF1* abundance of mRNA and both lesion severity and inflammatory infiltration support a link between histopathological changes and reduced expression of a growth factor.

## 1. Introduction

Equine endometrosis is a common cause of subfertility in mares and a major constraint on breeding efficiency worldwide (Katila and Ferreira-Dias, 2022; Schöniger and Schoon, 2020). In contrast to human endometriosis, which is defined by ectopic endometrial tissue (Zondervan et al., 2018), equine endometrosis is characterized by progressive fibrotic remodeling of the endometrial stroma and periglandular fibrosis with glandular alterations “fibrotic nests” (Kenney, 1993; Zdrojkowski et al., 2024). Disease severity is routinely assessed histopathologically, and advanced lesions are associated with reduced reproductive performance and limited therapeutic options because established fibrosis is largely irreversible (Amaral et al., 2021, 2020; Reghini et al., 2016). Although multiple molecular pathways have been implicated, the mechanisms linking specific histopathological features to local transcriptional changes in the equine endometrium remain incompletely defined (Jasiński et al., 2022a; Szóstek-Mioduchowska et al., 2022).

Fibrotic remodeling in endometrosis is closely related to fibroblast activation and differentiation toward a myofibroblast-like phenotype (Masseno, 2012). This cellular transition is commonly associated with increased alpha-smooth muscle actin (α-SMA) expression, encoded by *ACTA2*, and is influenced by transforming growth factor beta 1 (TGF-β1), which promotes extracellular matrix (ECM) deposition and stromal remodeling in a range of fibrotic tissues (Klose and Schoon, 2016; Skarzynski et al., 2020; Szóstek-Mioduchowska et al., 2019; Wong et al., 2023). Endometrial responses to profibrotic signaling occur within a hormonally regulated environment and altered expression or function of estrogen receptor α (ERα) encoded by *ESR1* and progesterone receptor (PR) encoded by *PGR* has been linked to endometrial remodeling and impaired cellular responsiveness in mares with endometrosis (Hoffmann et al., 2009; Jasiński et al., 2022a; Schöniger and Schoon, 2020). However, fibrotic progression is likely shaped not only by profibrotic signaling and steroid hormone regulation but also by reduced capacity for epithelial and stromal homeostasis and repair.

Insulin-like growth factor 1 (IGF-1) encoded by *IGF1* is a candidate mediator of endometrial maintenance because it supports cell survival, proliferation, and metabolic activity in endometrial epithelial and stromal compartments and interacts with local hormonal cues (Hernandez et al., 2020; Hung et al., 2013; Majchrzak-Baczmańska and Malinowski, 2016; Rutanen, 1998). In mares, components of the insulin-like growth factor system are involved in early pregnancy and endometrial function, but the relationship between *IGF1* mRNA expression and histopathological hallmarks of endometrosis has not been fully characterized (Gibson et al., 2022; Kalpokas et al., 2021). Tissue proliferative activity can be approximated by the proliferation marker Ki-67, encoded by *MKI67*, which provides an index of cellular cycling and regenerative potential in endometrial tissues (Köhne et al., 2024; Mambelli et al., 2014).

Most molecular studies of equine endometrosis have emphasized profibrotic pathways, including TGF-β1–associated signaling and matrix remodeling mediators (Rebordão et al., 2014; Szóstek-Mioduchowska et al., 2020; Wong et al., 2023). Fewer data address whether increasing severity of specific histopathological lesions is accompanied by reduced expression of growth- and proliferation-related transcripts that may reflect impaired regenerative capacity. The present study therefore examined whether markers of tissue maintenance and cellular proliferation vary with endometrosis severity and defined histopathological features hence the objective was to quantify *IGF1* and *MKI67* relative abundance of mRNA transcript in the equine endometrium in relation to endometrosis severity and selected histopathological features. Moreover, we also evaluated how these patterns relate to transcripts associated with fibrosis (*TGFB1* and *ACTA2*) and steroid hormone signaling (*ESR1* and *PGR)* to determine whether fibrotic remodeling and lesion severity are accompanied by changes consistent with reduced local proliferative and regenerative potential.

## 2. Materials and methods

### 2.1 Tissue collection

Endometrial tissue samples (n = 72) were collected post-mortem from mares of various breeds and ages at a commercial abattoir in Rawicz, Poland, in August 2023. Tissues were obtained as by-products of routine slaughter. No procedures were performed on live animals. From each uterus, two samples were collected from a standardized anatomical region to minimize site-related variability (uterine body approximately 5 cm from the cervix). One full-thickness sample of the uterine wall (approximately 2 × 2 cm) was immediately immersed in 4% buffered formaldehyde (Chempur, Piekary Śląskie, Poland) for histological processing. A second sample consisting of endometrium (approximately 1–2 g; myometrial contamination minimized by careful dissection) was placed into RNase-free cryogenic tubes (Biologix Group Ltd., Jinan, China), snap-frozen in liquid nitrogen (LN_2_), and subsequently transferred to −80 °C for long-term storage until RNA analysis.

The estrous cycle phase was assigned based on gross ovarian morphology. The luteal phase was defined by the presence of a corpus hemorrhagicum and/or corpus luteum, whereas the follicular phase was defined by a dominant follicle >20 mm in diameter in the absence of luteal structures (“Autumnal Transition Out of the Breeding Season,” 2011).

### 2.2 Histological processing and Kenney and Doig classification

Formalin-fixed uterine wall samples were processed using a standard paraffin histology workflow. After fixation in 4% buffered formaldehyde (Chempur, Piekary Śląskie, Poland) for a minimum of 24 h and a maximum of 48 h at room temperature, tissues were trimmed to include endometrium and underlying myometrium and placed in labeled cassettes (Simport Scientific, Beloeil, QC, Canada). Samples were dehydrated through a graded ethanol series using ethanol (Avantor Performance Materials Poland S.A., Gliwice, Poland) (70%, 80%, 90%, 96%, and 99.8–100% ethanol, 100% ethanol 30 min each, 100% ethanol 1 h), cleared in xylene (Chempur, Piekary Śląskie, Poland) (two changes, 1 h each), and infiltrated (three changes, 1 h each) and embedded in paraffin (Sigma-Aldrich, St. Louis, MO, USA). Paraffin blocks were cooled and stored at room temperature until sectioning. Serial sections were cut at 4 µm using a rotary microtome and mounted onto glass microscope slides (Superfrost, Thermo Fisher Scientific, Waltham, MA, USA).

For hematoxylin and eosin (H&E) staining, slides were deparaffinized in xylene (Chempur, Piekary Śląskie, Poland) (two changes, 3 min each), rehydrated through a descending ethanol series (Avantor Performance Materials Poland S.A., Gliwice, Poland) to distilled water, stained with hematoxylin (Harris hematoxylin, Chempur, Piekary Śląskie, Poland) for 2 min, differentiated in acid alcohol, and blued in running tap water. Afterwards, slides were counterstained with eosin (Eosin Y, Chempur, Piekary Śląskie, Poland) for 40 s, dehydrated in ascending ethanol, cleared in xylene, and mounted using DPX mountant (Sigma-Aldrich, St. Louis, MO, USA) and coverslips (Jiangsu Bernoy Lab Instrument Co., Ltd., Yancheng, China).

Stained slides were examined using a light microscope (BX43, Olympus Corporation, Tokyo, Japan) under standardized magnifications (overview at 40× and 100×; detailed assessment at 200× and 400×). Histopathological evaluation was performed by two independent observers with experience in equine uterine pathology (Ł.Z. & M.D). Observers were blinded to RT-qPCR results and to each other’s scores during the primary reading. Discrepancies in KD category or in additional scores were resolved by consensus review at joint session using the same microscope configuration (BX43, Olympus Corporation, Tokyo, Japan).

All samples were classified according to the Kenney and Doig (KD) system (Kenney, 1993, 1978) using predefined criteria for fibrosis severity and the proportion of affected glands, resulting in assignment to KD categories I, IIA, IIB, or III. KD assessment was completed for all n = 72 samples before any sample selection for molecular analyses to minimize the risk of interpretation bias based on gene expression data.

Selection of samples for RT-qPCR and definition of analytical groups. After screening all n = 72 samples, a subset (n = 47) was selected for RT-qPCR to represent the spectrum of endometrosis while limiting confounding by pronounced inflammatory lesions inconsistent with the study aim (endometrosis-associated remodeling rather than predominantly inflammation-driven changes). Selection was performed using an a priori workflow based on histology. First, all KD categories were assigned. Second, samples were reviewed for inflammatory activity using the overall inflammatory infiltration score defined in Table 1 (feature 11) and the cell-type–specific ordinal scales (features 12–15).

**Table 1.**
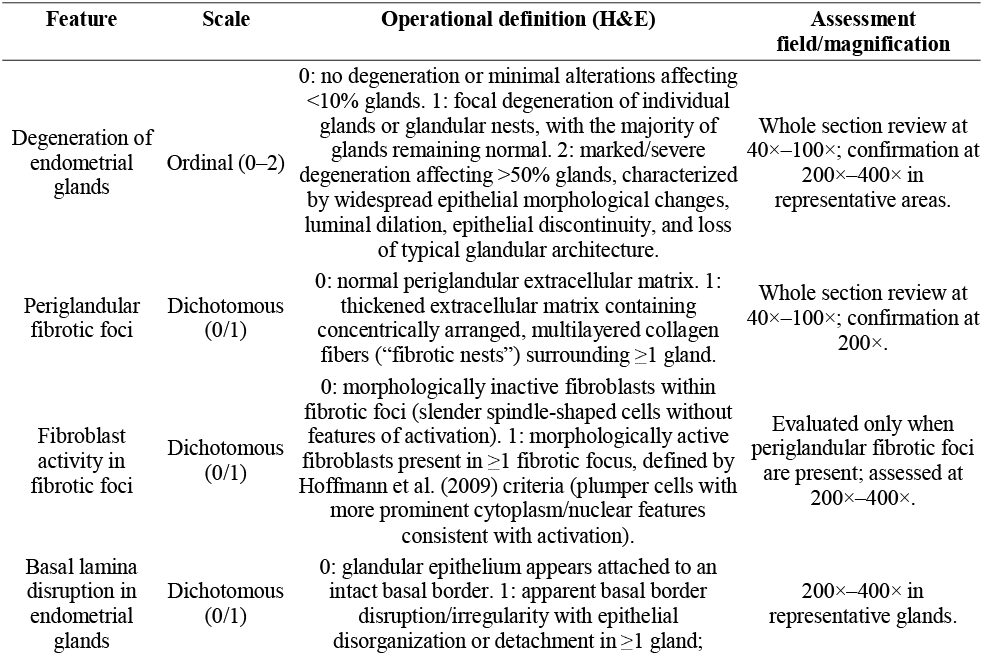
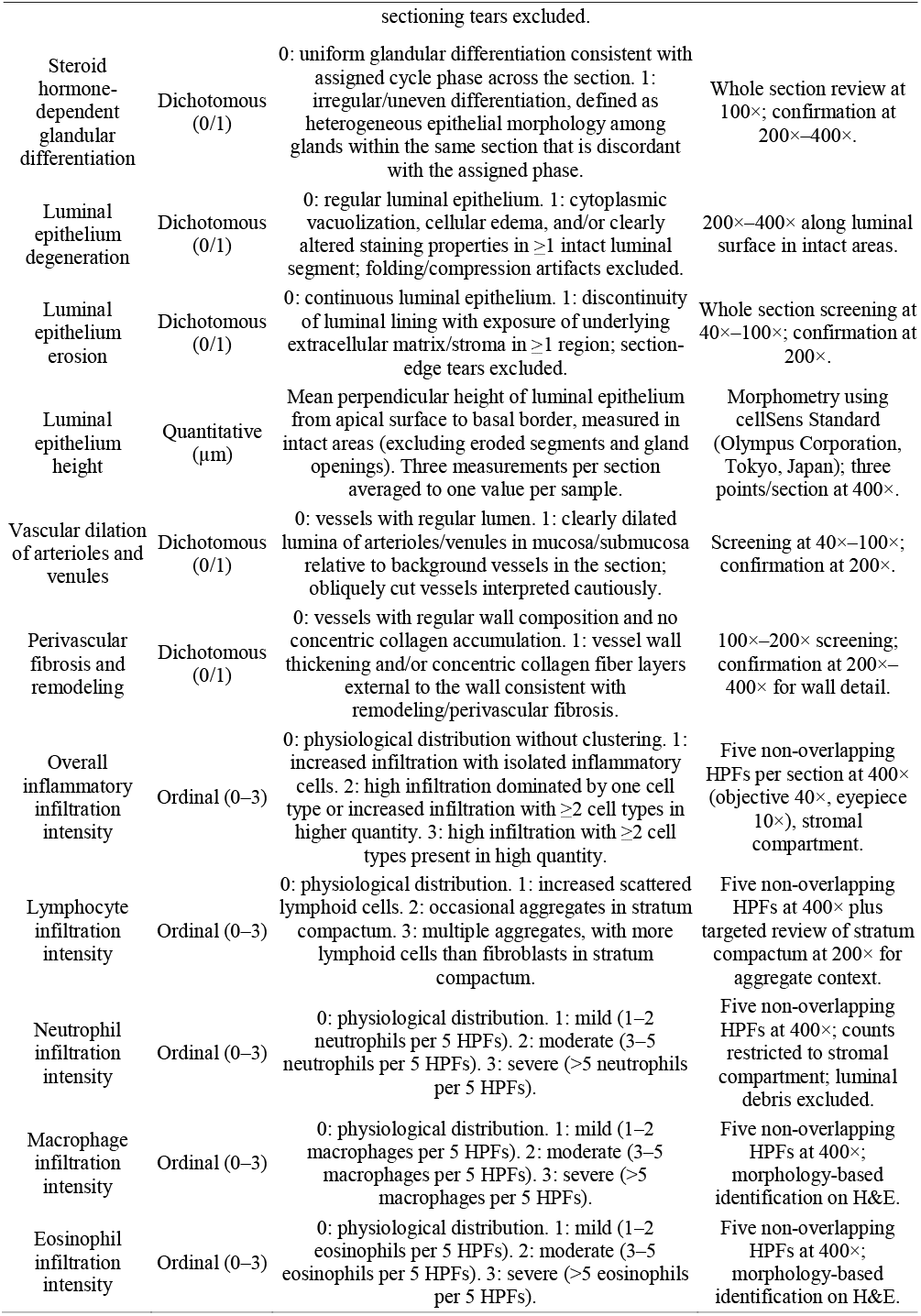
Histopathological features for the equine endometrial samples evaluation assessed additionally to Kenney and Doig (KD) categories (Kenney, 1993, 1978).

Samples were excluded from molecular analysis primarily if they met one or more of the following criteria: (i) absence of endometrosis-associated remodeling (KD I) combined with any evidence of active inflammation beyond physiological background (to maintain a clean control group), and/or (ii) technical factors that could compromise downstream analyses (e.g., advanced autolysis, extensive sectioning artifacts, insufficient tissue).

Because early lesions may not fulfill KD IIA criteria at the whole-section level, an additional “early/minor endometrosis” group (I+) was defined a priori to capture samples with focal endometrosis-compatible remodeling. I+ was operationalized as follows: (1) overall KD grading consistent with KD I (i.e., the sample did not meet KD IIA criteria for the extent of fibrosis/percentage of affected glands), and (2) concurrent presence of focal remodeling defined by at least one of the following additional features: periglandular fibrotic foci present (Table 1, feature 2 = 1) and/or glandular degeneration score 1 (Table 1, feature 1 = 1), and (3) absence of severe inflammation, defined as overall inflammatory infiltration score <3 (Table 1, feature 11 ≤2) and neutrophil infiltration intensity ≤1 (Table 1, feature 13 ≤1). This definition ensured that I+ captured early fibrotic/glandular remodeling while minimizing confounding by acute inflammatory activity. KD I samples nonclassified as I+ were retained as the control group.

The resulting molecular dataset comprised five analytical groups: KD I control (histologically normal endometrium without endometrosis and without endometritis; n = 10), I+ (early/minor endometrosis; n = 7), KD IIA (n = 11), KD IIB (n = 9), and KD III (n = 10).

### 2.3 Additional histopathological assessment

In addition to assigning a KD category, all endometrial samples included in the molecular dataset were independently evaluated using a structured panel of additional histopathological features (Table 1). Assessments were performed on hematoxylin and eosin (H&E)-stained sections prepared as described above and examined using a light microscope (BX43, Olympus Corporation, Tokyo, Japan). To ensure reproducibility, the same microscope configuration and magnification scheme were used for all slides. Each slide was first surveyed at low magnification (40× and 100×) to identify representative, well-preserved areas and to exclude regions affected by folds, tears, compression, edge artifacts, or advanced autolysis. Detailed scoring was subsequently performed at 200× and 400× magnification.

For dichotomous variables, a feature was recorded as present (score 1) when unequivocally identified in at least one representative region of the endometrium; otherwise, it was recorded as absent (score 0). For ordinal variables, scoring reflected the predominant pattern across the section rather than a focal maximum, unless the focal lesion was extensive enough to influence the overall tissue appearance. Inflammatory infiltration was assessed within the stromal compartment and scored using five non-overlapping high-power fields (HPFs) per section (400× total magnification; objective 40×, eyepiece 10×). HPFs were selected in representative areas of the stratum compactum and stratum spongiosum while avoiding glandular lumina, blood vessel lumina, and luminal debris to prevent systematic over- or underestimation of inflammatory cells.

Luminal epithelial height was quantified by calibrated morphometry using cellSens Standard software (Olympus Corporation, Tokyo, Japan). Measurements were performed at a fixed magnification (400×) in intact luminal epithelium, excluding eroded regions and gland openings. Three measurement points per section were recorded in µm and averaged to yield a single luminal epithelial height value per sample.

All additional features were evaluated independently by two observers under blinded conditions with respect to RT-qPCR results. Any discrepancies in feature scoring were resolved by a consensus review using the same microscope configuration (BX43, Olympus Corporation, Tokyo, Japan). Definitions and thresholds were applied consistently across all samples, and inflammatory-cell-type scoring followed established descriptions of equine endometrial inflammatory patterns (Jasiński et al., 2022b; Ricketts and Alonso, 1991).

### 2.4 RT-qPCR

Total RNA was isolated from frozen endometrial tissue using a phenol–guanidinium thiocyanate method. Briefly, approximately 100 mg of endometrium per sample was transferred into RNase-free, screw-cap homogenization tubes with ceramic beads (MagNA Lyser Green Beads, Roche Diagnostics, Basel, Switzerland). Tissue was immediately lysed in TRI Reagent (Sigma-Aldrich, St. Louis, MO, USA) and homogenized using a multi-tube bead shaker (MagNA Lyser, Roche Diagnostics, Basel, Switzerland) under standardized settings (frequency: 50-60 Hz, speed 7000 rpm, duration: 20 s; number of cycles: 2), with tubes kept cooled between cycles to limit RNA degradation.

RNA extraction was performed using TRI Reagent (Sigma-Aldrich, St. Louis, MO, USA) strictly according to the manufacturer’s protocol, including phase separation with chloroform (Chempur, Piekary Śląskie, Poland), RNA precipitation with isopropanol (Chempur, Piekary Śląskie, Poland), and washing with 75% ethanol prepared from molecular biology grade ethanol (Avantor Performance Materials Poland S.A., Gliwice, Poland) and RNase-free water (Chempur, Piekary Śląskie, Poland). RNA samples were stored at −80 °C until RT-qPCR analysis.

RNA concentration and purity were assessed by spectrophotometry using a NanoDrop Lite spectrophotometer (Thermo Fisher Scientific, Waltham, MA, USA). For each sample A260/280 ratio was recorded.

One-step RT-qPCR was performed on a LightCycler 480 II Real-time PCR system (Roche, Basel, Switzerland) using the TaqMan RNA-to-CT 1-Step Kit (Thermo Fisher Scientific, Waltham, MA, USA). Reactions were set up in 96-well plates (Thermo Fisher Scientific, Waltham, MA, USA) sealed with optical films (Thermo Fisher Scientific, Waltham, MA, USA) in a dedicated clean area to minimize contamination. Each sample was analyzed in technical triplicates. The total reaction volume was 18 µL per well.

Predesigned equine TaqMan Gene Expression Assays (Thermo Fisher Scientific, Waltham, MA, USA) were used for quantification of the following transcripts: Ec03470487_m1 (*SDHA*), Ec03210916_gH (*GAPDH*), Ec07039069_g1 (*MKI67*), Ec03468688_m1 (*IGF1*), Ec03468030_m1 (*TGFB1*), Ec07040967_m1 (*ACTA2*), Ec03467534_m1 (*ESR1*), and Ec03469694_s1 (*PGR*). All assays used a FAM reporter dye. Because commercial predesigned TaqMan assays, listed in Table 2, were used, primer and probe sequences and component concentrations are proprietary to the manufacturer and were therefore not reported; assays were applied at the recommended 1× final concentration.

**Table 2.** Details of the probes used for amplification of studied genes.

| Gene<br>(mRNA) | Gene name (protein) | RefSeq accession<br>number | TaqMan Assay<br>ID | Reporter<br>dye | Role |
| --- | --- | --- | --- | --- | --- |
| *SDHA* | Succinate dehydrogenase<br>complex flavoprotein subunit A<br>(SDHA) | XM_014734954.2 | Ec03470487_m1 | FAM | Reference<br>gene |
| *GAPDH* | Glyceraldehyde-3-phosphate<br>dehydrogenase (GAPDH) | NM_001163856.1 | Ec03210916_gH | FAM | Reference<br>gene |
| *MKI67* | Marker of proliferation Ki-67<br>(Ki-67) | XM_005602153.3 | Ec07039069_g1 | FAM | Target gene |
| *IGF1* | Insulin-like growth factor 1<br>(IGF-1) | NM_001082498.2 | Ec03468688_m1 | FAM | Target gene |
| *TGFB1* | Transforming growth factor beta<br>1 (TGF-β1) | NM_001081849.1 | Ec03468030_m1 | FAM | Target gene |
| *ACTA2* | Actin alpha 2, smooth muscle<br>(α-SMA) | XM_001503035.6 | Ec07040967_m1 | FAM | Target gene |
| *ESR1* | Estrogen receptor 1 (ERα) | NM_001081772.1 | Ec03467534_m1 | FAM | Target gene |
| *PGR* | Progesterone receptor (PR) | XM_023644695.1 | Ec03469694_s1 | FAM | Target gene |

Thermal cycling conditions were as follows: reverse transcription at 48 °C for 15 min, enzyme activation at 95 °C for 10 min, followed by 40 amplification cycles of denaturation at 95 °C for 15 s and annealing/extension at 60 °C for 60 s. No-template controls (NTCs; RNase-free water substituted for RNA) were included for each assay on every plate to monitor reagent contamination.

Quantification cycle (Cq/Ct) values were calculated using the second derivative maximum method in LightCycler 480 software version 1.5.1.62 (Roche, Basel, Switzerland). relative abundance of mRNA transcript was calculated using the comparative Ct method (2^−ΔΔCt^). The geometric mean of two reference genes (*SDHA* and *GAPDH*) was used for normalization (Kayis et al., 2011; Klein et al., 2011). The calibrator was defined as the control group (KD category I without histological signs of inflammation), using the mean ΔCt value of that group as the reference for ΔΔCt calculations. Results are presented as fold change in mRNA expression for categories I+, IIA, IIB, and III relative to healthy endometrium (KD I).

### 2.5 Statistical analysis

Statistical analyses were performed using IBM SPSS Statistics version 29 (Predictive Solutions, Warsaw, Poland). The analyses comprised: (i) comparisons of relative abundance of mRNA transcript among endometrosis categories (KD I, I+, IIA, IIB, and III); (ii) comparisons of relative abundance of mRNA transcript between groups defined by additional dichotomous histopathological features (e.g., absent vs present epithelial erosion, basal lamina disruption, or epithelial degeneration); and (iii) correlation analyses among transcripts and between relative abundance of mRNA transcripts and ordinal histopathological scores. Data visualization was performed in Python (Python Software Foundation, Wilmington, DE, USA).

Distributional assumptions for quantitative variables were assessed using the Shapiro–Wilk test. Because relative abundance of mRNA transcripts did not meet normality assumptions, nonparametric tests were applied. Differences in relative abundance of mRNA transcript among KD categories were tested using the Kruskal–Wallis test. When the omnibus test indicated a significant difference, it was followed by Dunn’s post-hoc test with Bonferroni correction for multiple pairwise comparisons. For comparisons involving dichotomous histopathological features, relative abundance of mRNA transcript was compared between groups using the Mann–Whitney U test.

Associations between continuous/ordinal variables were evaluated using Spearman’s rank correlation coefficient (*ρ*). Correlations were considered statistically significant at *P* < 0.05. Correlation strength was interpreted as weak (|*ρ*| = 0.10–0.29), moderate (|*ρ*| = 0.30–0.59), strong (|*ρ*| = 0.60–0.79), or very strong (|*ρ*| = 0.80–1.00) (Akoglu, 2018). All tests were two-tailed.

## 3. Results

### 3.1 Relative abundance of mRNA transcript in categories of endometrosis

Across endometrosis categories (Figure 1), *TGFB1* relative abundance of mRNA transcript was higher in category I+ than in category I (*P* = 0.024). No other pairwise differences in *TGFB1* relative abundance of mRNA transcript were detected. The relative abundance of mRNA transcript of *IGF1, MKI67, ACTA2, ESR1*, and *PGR* did not differ significantly among categories.

**Figure 1.**
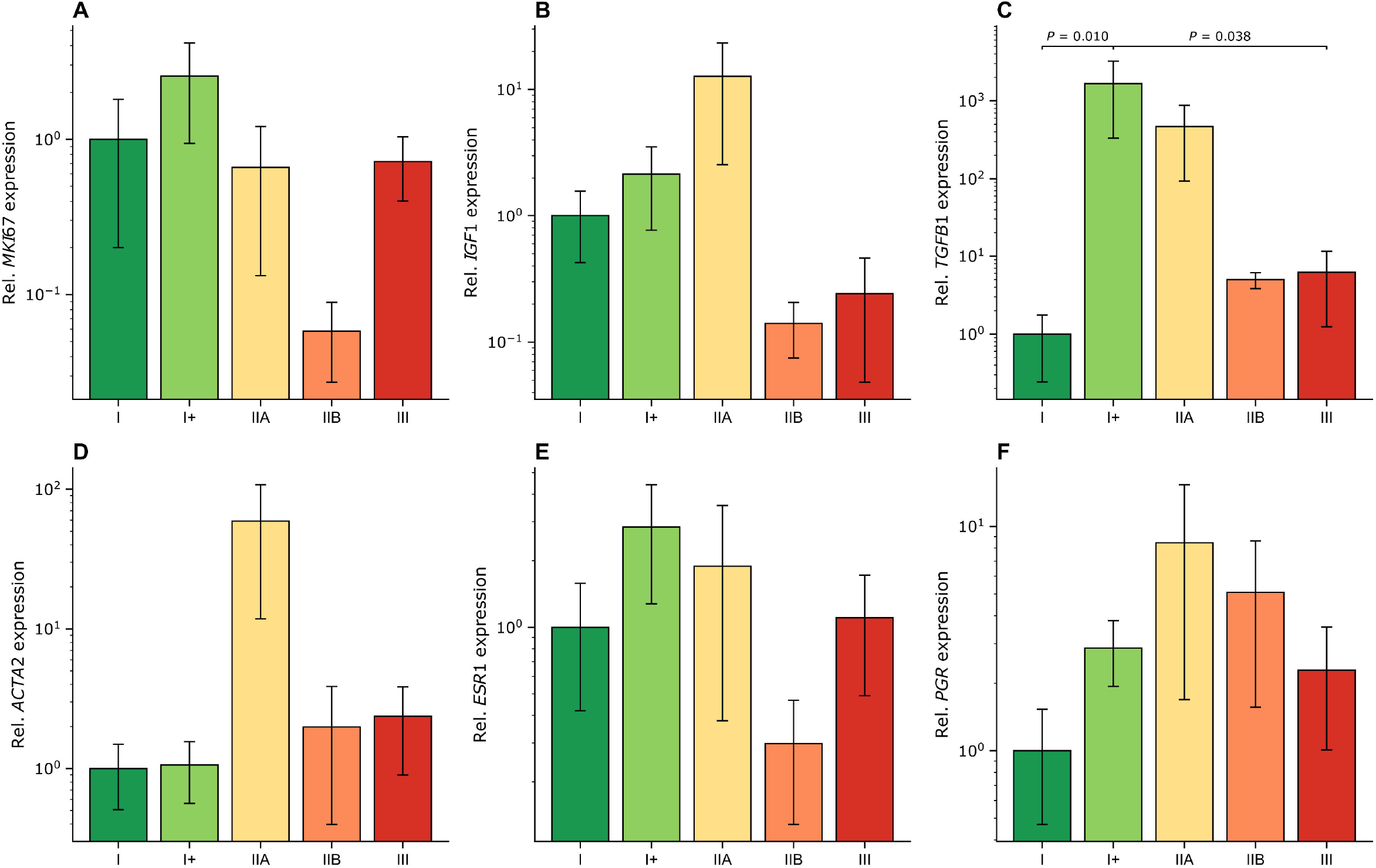
Relative abundance of mRNA transcript of *MKI67* (A), *IGF1* (B), *TGFB1* (C), *ACTA2* (D), *ESR1* (E), and *PGR* (F) across modified Kenney and Doig (KD) endometrosis categories (I, I+, IIA, IIB, III). Expression values were scaled so that the mean of KD category I equaled 1.0 for each transcript. Bars indicate the mean and error bars indicate the standard error of the mean (SEM). The y-axis is logarithmic.

### 3.2 Relative abundance of mRNA transcript in relation to additional histopathological features

When stratified by additional histopathological features (Figure 2), samples with luminal epithelial erosion showed lower *MKI67* relative abundance of mRNA transcript than samples with an intact luminal epithelium (*P* = 0.049). Samples with disruption of the glandular basal lamina showed higher *TGFB1* relative abundance of mRNA transcript than samples without basal lamina disruption (*P* = 0.020). Samples with luminal epithelial degeneration showed higher *ACTA2* relative abundance of mRNA transcript than samples without luminal epithelial degeneration (*P* = 0.044). No other comparisons reached statistical significance.

**Figure 2.**
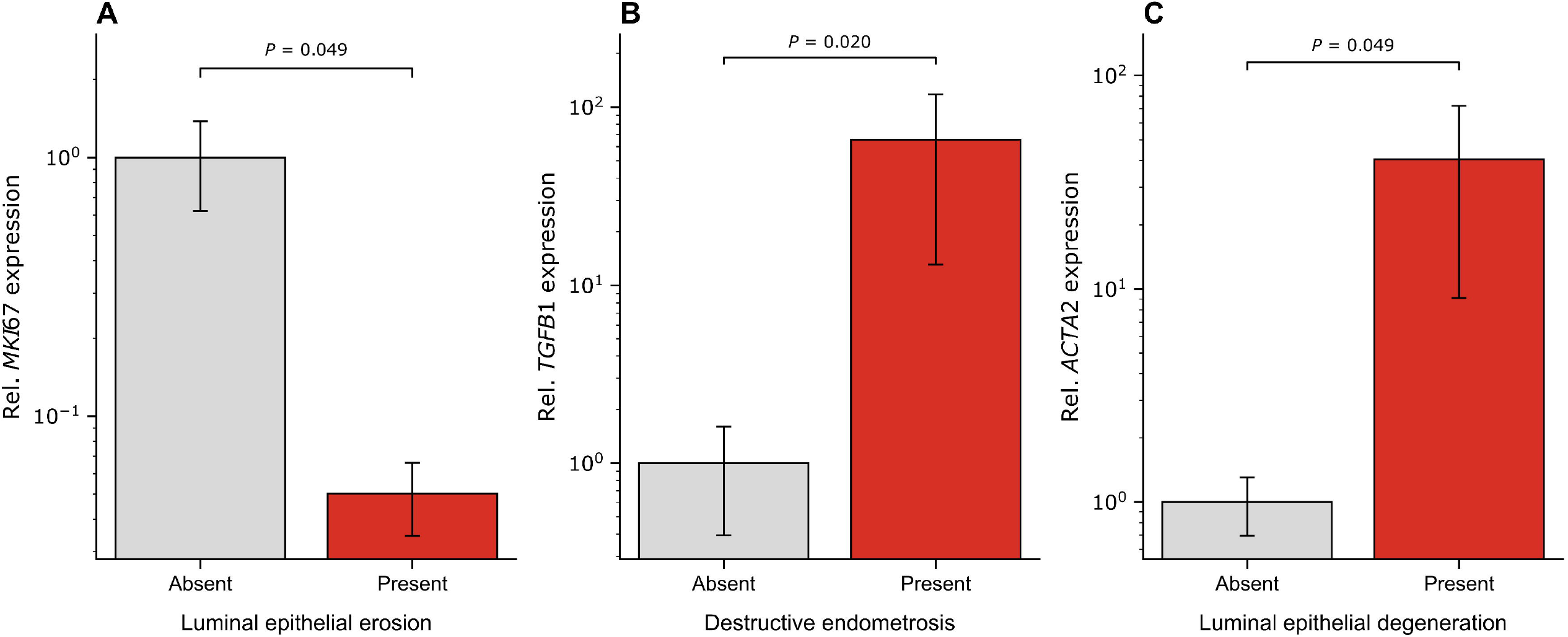
Relative abundance of mRNA transcript stratified by selected additional histopathological features. (A) *MKI67* expression by presence of luminal epithelial erosion (Absent vs Present). (B) *TGFB1* expression by presence of glandular basal lamina disruption (Absent vs Present). (C) *ACTA2* expression by presence of luminal epithelial degeneration (Absent vs Present). For each panel, expression values were scaled so that the mean of the Absent group equaled 1.0 for the corresponding transcript. Bars indicate the mean and error bars indicate SEM.

### 3.3 Correlations among relative abundance of mRNA transcripts and between relative abundance of mRNA transcripts and histopathological features

Significant positive correlations were identified among selected transcripts (Figure 3). *MKI67* relative abundance of mRNA transcript correlated positively with *ACTA2* (*ρ* = 0.557, *P* < 0.001), *ESR1* (*ρ* = 0.887, *P* < 0.001), and *PGR* (*ρ* = 0.429, *P* = 0.023). In addition, *ACTA2* correlated positively with *ESR1* (*ρ* = 0.585, *P* < 0.001) and *PGR* (*ρ* = 0.376, *P* = 0.024). *IGF1* relative abundance of mRNA transcript was not significantly correlated with *MKI67, TGFB1, ACTA2, ESR1*, or *PGR*.

**Figure 3.**
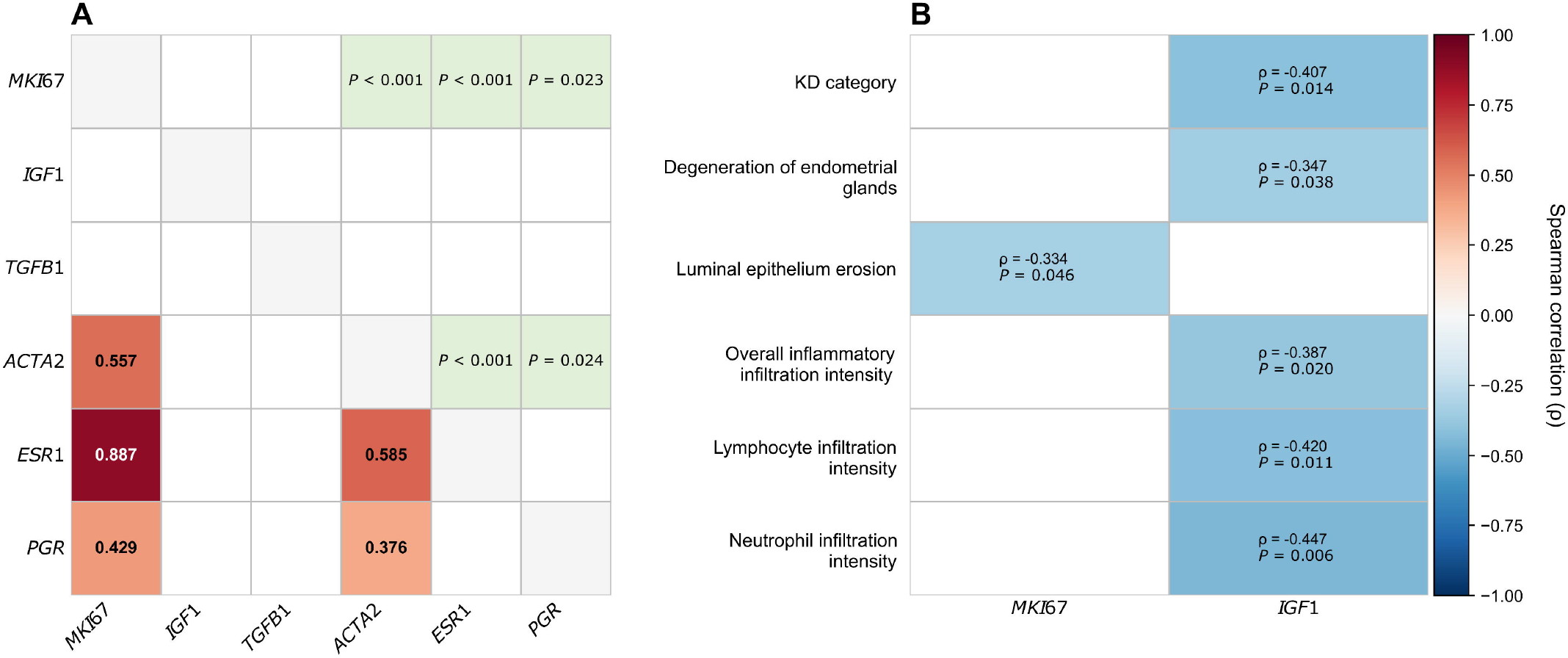
Spearman correlation analyses for relative abundance of mRNA transcript and histopathological features. (A) Pairwise correlations among *MKI67, IGF1, TGFB1, ACTA2, ESR1*, and *PGR*. The lower triangle shows Spearman’s rank correlation coefficients (*ρ*), and the upper triangle shows corresponding *P* values. Cells with statistically significant associations are highlighted in light green. Cell color in the lower triangle reflects the magnitude and direction of *ρ* (color bar). (B) Correlations between *MKI67* and *IGF1* relative abundance of mRNA transcript and selected histopathological features (KD category, degeneration of endometrial glands, overall inflammatory infiltration intensity, and inflammatory cell–type infiltration intensities). Each cell displays *ρ* and *P*; blank cells indicate no statistically significant correlation reported for the corresponding feature– transcript pair.

*MKI67* relative abundance of mRNA transcript was significantly correlated only with luminal epithelium erosion (*ρ* = −0.334, *P* = 0.046). In contrast, *IGF1* relative abundance of mRNA transcript showed negative correlations with selected histopathological endpoints. *IGF1* correlated negatively with KD category (*ρ* = −0.407, *P* = 0.014), degeneration of endometrial glands (*ρ* = −0.347, *P* = 0.038), overall inflammatory infiltration intensity (*ρ* = −0.387, *P* = 0.020), lymphocyte infiltration intensity (*ρ* = −0.420, *P* = 0.011), and neutrophil infiltration intensity (*ρ* = −0.447, *P* = 0.006). No significant correlations were observed between *IGF1* and macrophage or eosinophil infiltration intensity.

## 4. Discussion

Lower *MKI67* relative abundance of mRNA transcript in samples with luminal epithelial erosion, followed by moderate correlation, suggests reduced proliferative activity in endometrium exhibiting epithelial discontinuity. This finding is consistent with reports that Ki-67 expression decreases in epithelia with degenerative morphology and impaired cellular integrity (Dhaval, 2015). Because the present study quantified mRNA in whole endometrial tissue, the observed decrease cannot be attributed to a specific cell compartment; however, it is compatible with the interpretation that epithelial injury is accompanied by reduced local proliferative signaling and/or reduced representation of cycling cells in the sampled tissue. Prior work in mares has shown that Ki-67 expression in the luminal epithelium varies across the ovarian cycle and is typically lowest during the luteal phase (Gerstenberg et al., 1999). Therefore, cycle phase represents a potential confounder when interpreting proliferation-related transcripts in cross-sectional samples, and future studies assessing epithelial erosion should explicitly account for cycle phase and, where feasible, include endocrine measurements or more granular cycle staging (Schöniger et al., 2018). Conversely, it is also plausible that the presence of epithelial damage itself contributes to reduced proliferative activity, either through loss of viable epithelial cells or through local microenvironmental changes that suppress cell cycling (Jischa et al., 2008).

The positive correlation between *MKI67* and *ACTA2* relative abundance of mRNA transcript may reflect coordinated variation in cellular composition and/or activation states within endometrial tissue rather than direct proliferation of a single stromal cell type. In other fibrotic tissues, Ki-67 and α-SMA have been reported to co-vary, which has been interpreted as increased cellular activation within remodeled stroma (Kamala et al., 2022). In equine endometrium, both inflammatory infiltration and stromal remodeling can influence the relative abundance of activated stromal cells and immune cells, each contributing to bulk-tissue transcript levels (Aupperle et al., 2000; Hoffmann et al., 2009). Without compartment-specific localization (e.g., immunohistochemistry or in situ hybridization), the present data support an association between proliferation-related and cytoskeletal remodeling transcripts but do not establish which cells account for the signal. Analogously, elevated Ki-67 expression has been described in fibrotic stroma and immune cell populations in chronic liver disease in horses (Verhoef et al., 2018), supporting the concept that increased Ki-67 signal in fibrotic settings may partly reflect non-epithelial cell cycling or activation.

The very strong correlation between *MKI67* and *ESR1* relative abundance of mRNA transcript is in line with the hormonally regulated nature of endometrial proliferation and previously reported co-variation of Ki-67 and steroid receptor expression in equine endometrium (Aupperle et al., 2000). Estrogen-dominant conditions are associated with increased endometrial proliferation and glandular epithelial hyperplasia, consistent with higher Ki-67 expression (Hoffmann et al., 2009; Kilgenstein et al., 2015; Schöniger and Schoon, 2020). Accordingly, the observed relationship between *MKI67* and *ESR1* may reflect shared regulation by the hormonal milieu, and this should be considered when interpreting *MKI67* as an indicator of regenerative capacity in samples collected at heterogeneous cycle stages.

The negative correlation between *IGF1* relative abundance of mRNA transcript and KD category suggests that lower *IGF1* expression is associated with more advanced histopathological severity. Because IGF-1 supports cell survival, proliferation, and metabolic activity in multiple tissues, reduced *IGF1* relative abundance of mRNA transcript could be consistent with diminished tissue maintenance signaling in endometrium with progressive fibrotic remodeling (Hernandez et al., 2020; Hung et al., 2013; Majchrzak-Baczmańska and Malinowski, 2016; Rutanen, 1998). In mares, increased glandular development and endometrial activity during pregnancy have been linked to the insulin-like growth factor system, including IGF-1, even in endometria affected by endometrosis (Bracher et al., 1996; Kalpokas et al., 2021). The present study extends this context by indicating that, outside pregnancy, *IGF1* transcript levels are inversely associated with fibrosis severity and glandular degeneration scores. These findings do not establish causality; they may reflect reduced production of *IGF1* mRNA by specific endometrial compartments, altered cellular composition in advanced lesions, or suppression of *IGF1* relative abundance of mRNA transcript under pathological conditions.

*IGF1* relative abundance of mRNA transcript also correlated negatively with overall inflammatory infiltration, lymphocytic infiltration, and neutrophilic infiltration, whereas no comparable associations were detected for macrophage or eosinophil scores. This pattern is consistent with reports that inflammatory states can be accompanied by lower IGF-1 expression or altered insulin-like growth factor signaling in other models, potentially mediated by pro-inflammatory cytokines and changes in IGF-binding proteins (Grzędzicka et al., 2023; Heemskerk et al., 1999; Nederlof et al., 2022; Reidel et al., 2022). In horses, *IGF1* mRNA has been proposed as a negative acute-phase marker in certain inflammatory settings (Grzędzicka et al., 2023). In experimental models, IGF-1 deficiency has been associated with increased lymphocytic inflammation, whereas exogenous IGF-1 reduced inflammatory severity (Casellas et al., 2006; Morales-Garza et al., 2017). Although the equine endometrium differs substantially from systemic inflammatory models and from pregnancy-associated immune modulation (Camacho et al., 2020; Schöniger and Schoon, 2020), the inverse relationship between *IGF1* mRNA and inflammatory scores observed here supports the concept that reduced *IGF1* relative abundance of mRNA transcript accompanies a more inflammatory histopathological phenotype in non-pregnant tissue.

The association between lower *IGF1* relative abundance of mRNA transcript and neutrophilic infiltration should be interpreted cautiously. Experimental studies suggest that IGF-1 can modulate neutrophil phenotype and may reduce the propensity to form neutrophil extracellular traps (NETs) in certain contexts (Nederlof et al., 2022; Reidel et al., 2022). However, NET formation and neutrophil phenotypes were not directly assessed in this study, and bulk-tissue mRNA measurements cannot resolve whether local IGF-1 signaling affects neutrophil function within the equine endometrium. Consequently, the present data support an association between neutrophil burden and lower *IGF1* relative abundance of mRNA transcript but do not justify mechanistic inference regarding NETs in endometrosis-affected tissues.

Higher *TGFB1* relative abundance of mRNA transcript in category I+ compared with categories I and III suggests that profibrotic signaling at the transcript level may be particularly evident in early, subtle lesions captured by the I+ definition. Because TGF-β1 is produced as a latent complex and its biological activity is strongly regulated post-translationally, mRNA abundance does not necessarily reflect active TGF-β1 signaling within tissue (Kim et al., 2018). Nonetheless, increased *TGFB1* transcription in early lesions is consistent with a role for TGF-β1–associated pathways during the initiation or early amplification of stromal remodeling (Skarzynski et al., 2020; Szóstek-Mioduchowska et al., 2019; Wong et al., 2023). Nevertheless, results suggest that additional subdivision encompassing minor, initial lesions associated with endometrosis (category I+) may be a useful addition in studies regarding pathogenesis of endometrosis.

The association between higher *TGFB1* relative abundance of mRNA transcript and glandular basal lamina disruption is compatible with the concept that basement membrane injury may accompany early remodeling and can coincide with increased cytokine signaling (Klose and Schoon, 2016). Prior work indicates that macrophages can contribute to TGF-β1 production in equine endometrium under inflammatory stimulation (Skarzynski et al., 2020), and MCP-1–related pathways, which may be influenced by TGF-β1, have been linked to macrophage infiltration in equine endometrium (Arici et al., 1999; Cheng et al., 2005; Jasiński et al., 2022b). In the present study, macrophage infiltration was assessed morphologically on H&E sections, which may not capture macrophage activation states or functional polarization. Therefore, the lack of an association between macrophage score and *TGFB1* relative abundance of mRNA transcript does not exclude macrophage contribution to cytokine signaling; it may reflect that activation state rather than cell counts is more closely related to local cytokine transcription (Skarzynski et al., 2020; Szóstek-Mioduchowska et al., 2019).

Higher *ACTA2* relative abundance of mRNA transcript in samples with luminal epithelial degeneration suggests that epithelial injury is accompanied by increased expression of a transcript often associated with myofibroblast differentiation and cytoskeletal remodeling. However, because *ACTA2* mRNA was measured in homogenized tissue, the data do not localize the source of *ACTA2* relative abundance of mRNA transcript to specific stromal cells, vascular smooth muscle cells, or other compartments. Future studies using α-SMA immunolabeling or spatial methods would be required to determine whether increased *ACTA2* mRNA reflects myofibroblast accumulation specifically within fibrotic foci and how this relates to epithelial degeneration.

Present study has several limitations that should be considered when interpreting the findings. The design was cross-sectional and based on post-mortem abattoir material collected within a single month, which limits inference regarding temporal progression and may introduce selection bias. Clinical reproductive history, parity, and detailed health status were not available for all mares, which may have contributed unmeasured variability in both histopathological phenotype and gene expression. Estrous cycle phase was assigned from gross ovarian morphology rather than endocrine measurements. Because proliferation- and steroid receptor-related transcripts vary across the cycle, residual confounding is possible, particularly for *MKI67, ESR1*, and *PGR* (Gerstenberg et al., 1999). Gene expression was quantified as mRNA in whole-tissue homogenates, and bulk measurements are influenced by tissue composition. Fibrosis, epithelial injury, and immune cell infiltration can alter the proportional contribution of individual cell types to the transcript pool. Therefore, the present results cannot localize changes to specific compartments or distinguish reduced expression within a cell type from reduced abundance of that cell type. In addition, transcriptional findings were not verified at the protein level. Ki-67 and α-SMA are typically interpreted using immunohistochemistry, and TGF-β1 activity is regulated post-translationally, including latent and active forms, so functional signaling cannot be inferred directly from mRNA abundance (Kim et al., 2018). Histopathological scoring, although standardized, remained semi-quantitative and potentially subject to inter-observer variability. The observed associations should therefore be interpreted as hypothesis-generating signals requiring confirmation in independent cohorts or in models incorporating adjustment for multiple testing and relevant covariates.

## 5. Conclusions

The presence of luminal epithelial erosion was associated with lower *MKI67* relative abundance of mRNA transcript, consistent with reduced proliferative activity in samples exhibiting epithelial damage. *IGF1* relative abundance of mRNA transcript correlated negatively with KD category, glandular degeneration, and the intensity of overall inflammation, lymphocytic infiltration, and neutrophilic infiltration, indicating that lower *IGF1* relative abundance of mRNA transcript is associated with more advanced histopathological lesions and a higher inflammatory cell burden. In addition, *TGFB1* relative abundance of mRNA transcript was higher in category I+ than in categories I and III, supporting an association between early endometrosis-compatible remodeling and increased profibrotic signaling at the transcript level.

## Supporting information

Supplementary materials - quartile values

## CRediT authorship contribution statement

**Łukasz Zdrojkowski:** Conceptualization, Formal analysis, Investigation, Data Curation, Writing - Original Draft, Funding acquisition. **Anna Niwi**ń**ska:** Formal analysis, Investigation. **Ewa Kautz:** Methodology, Investigation. **Dawid Tobolski:** Formal analysis, Data Curation, Visualization Writing - Review & Editing. **Magdalena Fajkowska:** Methodology, Validation, Writing - Review & Editing. **Małgorzata Rzepkowska:** Methodology, Validation, Writing - Review & Editing. **Tomasz Jasi**ń**ski:** Validation, Writing - Original Draft, Writing - Review & Editing. **Małgorzata Domino:** Investigation, Validation, Resources, Project administration, Funding acquisition. **Bartosz Pawli**ń**ski:** Conceptualization, Resources, Supervision.

## Data and availability statement

The data generated during this study have been deposited in the Zenodo repository at 10.5281/zenodo.20326518. Further information is available from the corresponding author on reasonable request.

