## Supplementary materials - quartile values for "Evaluation of *IGF1* and *MKI67* relative abundance of mRNA transcript in relation to histopathological features of equine endometrosis"

| Gene | KD category | Q1 | Me | Q3 |
| --- | --- | --- | --- | --- |
| *MKI67* | I | ,077091115073787 | ,236674433599137 | 9,701159465086755 |
|  | I+ | ,761765624399980 | 7,811207550629079 | 38,729143375968480 |
|  | IIA | ,490846384655610 | 1,446874509644896 | 9,639906944370829 |
|  | IIB | ,092694581180107 | ,232910086748650 | ,719434452011227 |
|  | III | ,728534112783647 | 3,082937205192350 | 16,820343617707040 |
| *ESR1* | I | ,159833680865064 | 1,405698435472364 | 7,694818095026737 |
|  | I+ | ,394747854231366 | 3,611232569984251 | 35,599304339145390 |
|  | IIA | ,051888384028465 | ,986867375153423 | 2,841837110225210 |
|  | IIB | ,250779864846500 | ,557591338329416 | 1,838887824801184 |
|  | III | ,684228335584983 | 2,307318952158286 | 4,883245064400443 |
| *PGR* | I | ,421567611161956 | 1,566211384008630 | 20,792039710734166 |
|  | I+ | 4,457214050085860 | 15,802080930788570 | 38,471730784380450 |
|  | IIA | 1,558414956791264 | 10,400054999216735 | 32,401029734571320 |
|  | IIB | ,008139796064044 | 15,173791683782780 | 66,627518031580650 |
|  | III | ,653136344668116 | 7,410499838906612 | 23,625078964503130 |
| *TGFB1* | I | ,052743849793846 | ,268019778702527 | 1,921072477637569 |
|  | I+ | 7,419988789899723 | 93,166786897450150 | 1909,928490607495200 |
|  | IIA | 12,178172036580277 | 66,387671615319290 | 969,493793599069100 |
|  | IIB | 2,867334002157820 | 3,732881922653150 | . |
|  | III | ,358389309258368 | ,945122020882782 | 12,634819825055814 |
| *ACTA2* | I | ,058095455889306 | 2,109695591762169 | 19,078103048307746 |
|  | I+ | ,844325538039913 | 3,577264215865210 | 16,979384805094780 |
|  | IIA | 2,121776216906150 | 21,749726914057124 | 221,912193118677300 |
|  | IIB | ,026393383227430 | ,692405392101220 | 2,954708022212548 |
|  | III | 1,328617027019754 | 2,868194105213445 | 24,390091080854575 |
| *IGF1* | I | ,318625921506277 | ,451866321967345 | 10,379487088528090 |
|  | I+ | ,754143554369647 | 1,710176269542400 | 11,077286496339692 |
|  | IIA | ,008558518031048 | ,158143090270271 | 31,911422060150507 |
|  | IIB | ,031666392670513 | ,355619722756008 | ,944569760834355 |
|  | III | ,007243547728959 | ,036599175308442 | ,339162861537420 |

| Gene | Fibroblast activity | Q1 | Me | Q3 |
| --- | --- | --- | --- | --- |
| *MKI67* | 0 | ,212893379749765 | ,740600038205604 | 1,871304088929927 |
|  | 1 | ,231485926721400 | 4,146654080201220 | 17,788276192962194 |
| *ESR1* | 0 | ,160399717904060 | 1,198242815802254 | 3,389952689563915 |
|  | 1 | ,291953952815172 | 1,512125474045565 | 11,616418795416909 |
| *PGR* | 0 | ,530054634823828 | 8,004761611198862 | 27,627787766405350 |
|  | 1 | 1,524895581499470 | 10,400054999216735 | 28,336591966671723 |
| *TGFB1* | 0 | ,792849265494994 | 4,598429843148480 | 47,483594253543740 |
|  | 1 | ,172750567695673 | 5,559141997688350 | 117,570082499971050 |
| *ACTA2* | 0 | ,486239218594805 | 2,538214723906008 | 17,017758983267942 |
|  | 1 | ,315350017284965 | 11,655181008600861 | 37,027171354095000 |
| *IGF1* | 0 | ,036599175308442 | ,339162861537420 | 10,379487088528090 |
|  | 1 | ,091813448507637 | ,451866321967345 | 1,378143687137240 |

| Gene | Destructive endometrosis | Q1 | Me | Q3 |
| --- | --- | --- | --- | --- |
| *MKI67* | 0 | ,228637606666901 | ,764255393930887 | 7,478031394529106 |
|  | 1 | ,563089721604770 | 1,734624557259407 | 8,931367380343290 |
| *ESR1* | 0 | ,250779864846500 | ,817217416538734 | 3,777638559219151 |
|  | 1 | ,469356109489318 | 2,099717902987994 | 4,725463336730122 |
| *PGR* | 0 | ,500978930810429 | 11,333042758942707 | 25,617260265439235 |
|  | 1 | 1,432207854654648 | 6,303420472642025 | 33,551106079209810 |
| *TGFB1* | 0 | ,356473140356441 | 1,762061082250353 | 21,481760574001427 |
|  | 1 | 4,165655882900815 | 117,570082499971050 | 2718,562744230999700 |
| *ACTA2* | 0 | ,124573130093585 | 2,107164575511497 | 19,555706870662554 |
|  | 1 | 2,121776216906150 | 7,391402358411920 | 32,815860233475930 |
| *IGF1* | 0 | ,028262648885863 | ,586795364325973 | 4,949168769315167 |
|  | 1 | ,048930379934637 | ,390731685011644 | 1,334910781186931 |

| Gene | Luminal epithelium continuity | Q1 | Me | Q3 |
| --- | --- | --- | --- | --- |
| *MKI67* | 0 | ,197856210356514 | ,584638807035348 | ,830008867742472 |
|  | 1 | ,276101746950947 | 1,734624557259407 | 14,576379136271221 |
| *ESR1* | 0 | ,250779864846500 | ,817217416538734 | 1,838887824801184 |
|  | 1 | ,254037980615910 | 1,641757863607203 | 5,456873303722377 |
| *PGR* | 0 | 1,511123647329750 | 20,792039710734166 | 31,910897844152930 |
|  | 1 | ,474508490927605 | 5,541533368026351 | 25,950312346205866 |
| *TGFB1* | 0 | 2,094394235240245 | 3,732881922653150 | 380,025763851086250 |
|  | 1 | ,356473140356441 | 6,799706525829265 | 73,074799695869610 |
| *ACTA2* | 0 | ,248980196808458 | 2,853585266925654 | 20,996064929449810 |
|  | 1 | ,379313104831280 | 3,131118854801242 | 21,749726914057124 |
| *IGF1* | 0 | ,023725377865616 | ,192951596615909 | 7,106057660210487 |
|  | 1 | ,042202293499382 | ,451866321967345 | 3,006978470722900 |

| Gene | Luminal epithelium degeneration | Q1 | Me | Q3 |
| --- | --- | --- | --- | --- |
| *MKI67* | 0 | ,113167257323191 | ,766745163461794 | 7,811207550629079 |
|  | 1 | ,416969241614452 | 1,161732114936205 | 8,398605205336107 |
| *ESR1* | 0 | ,110312314672946 | 1,315005156041931 | 3,002266819908678 |
|  | 1 | ,346530162757600 | 1,728720168709975 | 4,883245064400443 |
| *PGR* | 0 | ,083444384883806 | 4,999373709056106 | 26,190652085083432 |
|  | 1 | 1,390892052145488 | 14,102655437608547 | 29,284460181275435 |
| *TGFB1* | 0 | ,409697964259120 | 5,559141997688350 | 36,485157772053020 |
|  | 1 | ,503209294670577 | 4,598429843148480 | 117,570082499971050 |
| *ACTA2* | 0 | ,062038559236332 | 2,113204888083487 | 11,157754091705740 |
|  | 1 | ,844325538039913 | 3,631754651166066 | 46,391964839094996 |
| *IGF1* | 0 | ,055658466369891 | ,451866321967345 | 10,728386792433891 |
|  | 1 | ,025483806745003 | ,339162861537420 | 1,528499856792692 |

| Gene | Perivascular fibrosis | Q1 | Me | Q3 |
| --- | --- | --- | --- | --- |
| *MKI67* | 0 | ,173038672035920 | ,952853999385773 | 9,064957517536055 |
|  | 1 | ,246135420558623 | ,719434452011227 | 6,478502926229186 |
| *ESR1* | 0 | ,260828861352018 | 1,198242815802254 | 3,694435564601701 |
|  | 1 | ,233631301718308 | 1,925802008806826 | 5,581227345955705 |
| *PGR* | 0 | ,474508490927605 | 5,541533368026351 | 24,367054653651940 |
|  | 1 | 1,030197266592703 | 15,802080930788570 | 31,910897844152930 |
| *TGFB1* | 0 | ,406902855059494 | 5,559141997688350 | 98,666005138195500 |
|  | 1 | 1,097394776270570 | 4,598429843148480 | 85,291748977094840 |
| *ACTA2* | 0 | ,240632575150295 | 2,046054660151647 | 18,523422935508663 |
|  | 1 | 2,121776216906150 | 7,391402358411920 | 46,391964839094996 |
| *IGF1* | 0 | ,192951596615909 | ,754143554369647 | 10,379487088528090 |
|  | 1 | ,031041491026723 | ,143882081376972 | ,782199476814093 |

| Gene | Vascular ectasia | Q1 | Me | Q3 |
| --- | --- | --- | --- | --- |
| *MKI67* | 0 | ,292397847054712 | 1,161732114936205 | 7,811207550629079 |
|  | 1 | ,096670931128060 | ,761765624399980 | 7,544027276243551 |
| *ESR1* | 0 | ,265818890356037 | 1,618552512618765 | 5,938562676653789 |
|  | 1 | ,221207281691152 | 1,007730116170494 | 3,303821027318609 |
| *PGR* | 0 | ,479719019188754 | 5,541533368026351 | 24,367054653651940 |
|  | 1 | 1,030197266592703 | 15,802080930788570 | 52,195601452177730 |
| *TGFB1* | 0 | ,404107745859868 | 2,867334002157820 | 36,485157772053020 |
|  | 1 | 2,695639554321737 | 149,848416022847260 | 725,138306057390900 |
| *ACTA2* | 0 | ,128152899149696 | 3,168955809945300 | 19,733097801336513 |
|  | 1 | ,935383641198588 | 2,208299109137557 | 37,172207817804450 |
| *IGF1* | 0 | ,098203970176449 | ,451866321967345 | 10,379487088528090 |
|  | 1 | ,009047564075875 | ,339162861537420 | ,900127329550863 |

| Gene | Foci of periglandular fibrosis | Q1 | Me | Q3 |
| --- | --- | --- | --- | --- |
| *MKI67* | 0 | ,241404927078880 | 1,249381033907691 | 10,908855004739374 |
|  | 1 | ,197856210356514 | ,836144225698500 | 5,196636032590351 |
| *ESR1* | 0 | ,159833680865064 | 1,458122103077183 | 4,455162073018154 |
|  | 1 | ,283301877884692 | 1,007730116170494 | 4,345100082521896 |
| *PGR* | 0 | ,537870427215552 | 9,111840977463448 | 28,909983786315920 |
|  | 1 | ,778823318642979 | 6,897682244934275 | 27,512698734027698 |
| *TGFB1* | 0 | ,405505300459681 | 3,992934692959243 | 121,507601460148690 |
|  | 1 | 1,311470449660700 | 10,539529008899688 | 75,839710296207070 |
| *ACTA2* | 0 | ,728072429737755 | 3,354191535333226 | 19,051549592482402 |
|  | 1 | ,128152899149696 | 2,311964659014290 | 33,459273722564090 |
| *IGF1* | 0 | ,082802771459447 | ,586795364325973 | 4,850105625174198 |
|  | 1 | ,009949572249332 | ,266057229076665 | 2,545516276472666 |

| Gene | Glandular degeneration | Q1 | Me | Q3 |
| --- | --- | --- | --- | --- |
| *MKI67* | 0 | ,179651338940907 | 1,737232210165227 | 11,499002525035719 |
|  | 1 | ,243509546743416 | ,710252990355699 | 5,427257125977896 |
|  | 2 | ,232910086748650 | ,952853999385773 | 4,769347068044072 |
| *ESR1* | 0 | ,239680690108954 | 1,431910269274773 | 7,352273406616901 |
|  | 1 | ,204367404449526 | 1,728720168709975 | 4,315783541097698 |
|  | 2 | ,414234598093476 | ,817217416538734 | 3,457920161472248 |
| *PGR* | 0 | ,450643315175355 | 4,633929319458730 | 26,880497767420480 |
|  | 1 | 1,652997575714291 | 14,954821482601123 | 27,064557620883413 |
|  | 2 | ,404612349329392 | 3,369677983471239 | 44,445062926138505 |
| *TGFB1* | 0 | 1,279329599887661 | 8,040271053970180 | 167,948609832284200 |
|  | 1 | ,406902855059494 | 1,097394776270570 | 66,387671615319290 |
|  | 2 | 4,598429843148480 | 10,539529008899688 | . |
| *ACTA2* | 0 | ,379313104831280 | 2,538214723906008 | 16,979384805094780 |
|  | 1 | ,211090943626278 | 14,719210985834968 | 39,925619280829544 |
|  | 2 | ,696388797909550 | 2,104633559260825 | 12,223767759926512 |
| *IGF1* | 0 | ,318625921506277 | ,762084936408111 | 11,077286496339692 |
|  | 1 | ,009875192592430 | ,192951596615909 | 5,596565535512590 |
|  | 2 | ,009101893660439 | ,204138750388910 | ,576197485550808 |

| Gene | Inflammation | Q1 | Me | Q3 |
| --- | --- | --- | --- | --- |
| *MKI67* | 0 | ,149932073434987 | ,766745163461794 | 60,597915732115080 |
|  | 1 | ,100979710196858 | ,703178657619294 | 1,794108151718337 |
|  | 2 | ,230061766694151 | ,719434452011227 | 15,293165666321343 |
|  | 3 | ,787739961303008 | 5,623924997136630 | 25,519359559793802 |
| *ESR1* | 0 | ,087353685724726 | 1,386563629559557 | 10,308189053592880 |
|  | 1 | ,181065895895004 | ,374965045959723 | 2,520977690858273 |
|  | 2 | ,250779864846500 | 1,405698435472364 | 5,299091576052057 |
|  | 3 | 1,866333283982803 | 4,155946241420981 | 16,050484733483522 |
| *PGR* | 0 | 1,700288885175805 | 5,541533368026351 | 25,284208184672604 |
|  | 1 | ,051882318453140 | 1,538667515669190 | 20,975986977078954 |
|  | 2 | 5,709158700349775 | 31,910897844152930 | 82,047558172095240 |
|  | 3 | ,399767568798628 | 7,840631800693917 | 18,505664825026210 |
| *TGFB1* | 0 | ,309667775030750 | 3,992934692959243 | 17,901101853863572 |
|  | 1 | 1,638515983208470 | 36,485157772053020 | 725,138306057390900 |
|  | 2 | 1,097394776270570 | 16,480628174650896 | 149,848416022847260 |
|  | 3 | ,213569323846160 | ,503209294670577 | . |
| *ACTA2* | 0 | ,379313104831280 | 2,538214723906008 | 18,423108295278983 |
|  | 1 | ,158882614219655 | 2,645366013425725 | 19,505915065051810 |
|  | 2 | ,301766587782320 | 3,604509433515638 | 20,576328475903118 |
|  | 3 | 1,328617027019754 | 17,781953640912460 | 68,672463656517340 |
| *IGF1* | 0 | ,607534222650326 | 6,044831679035246 | 16,471992232430438 |
|  | 1 | ,098203970176449 | ,721724406684600 | 11,077286496339692 |
|  | 2 | ,021825750164450 | ,113628864755335 | ,576197485550808 |
|  | 3 | ,005548190551173 | ,339162861537420 | 3,321338320122335 |

| Gene | Lymphocytes | Q1 | Me | Q3 |
| --- | --- | --- | --- | --- |
| *MKI67* | 0 | ,246135420558623 | ,506440292010209 | . |
|  | 1 | ,088545899710504 | ,719434452011227 | 4,813006392401166 |
|  | 2 | ,354683544334582 | 1,057293057160989 | 4,145390232339061 |
|  | 3 | 3,285598753704647 | 20,267623848199660 | 29,922620831852030 |
| *ESR1* | 0 | ,000967704381612 | 1,458122103077183 | . |
|  | 1 | ,116149012839097 | ,394747854231366 | 2,360547981174805 |
|  | 2 | ,580314352906336 | 1,838887824801184 | 4,916672749837434 |
|  | 3 | ,188090728857451 | 4,853928522976244 | 34,359947712388305 |
| *PGR* | 0 | 2,660600005888441 | 13,166786539380258 | 24,161166066187995 |
|  | 1 | ,051882318453140 | 1,052040905612091 | 25,071048974842430 |
|  | 2 | 1,390892052145488 | 12,185208803935282 | 41,902698451329770 |
|  | 3 | 4,284274191220605 | 15,226573632791850 | 40,474201923828154 |
| *TGFB1* | 0 | ,404107745859868 | 2,426727388230136 | . |
|  | 1 | 1,456880933981460 | 6,799706525829265 | 148,657892174304150 |
|  | 2 | 1,097394776270570 | 16,480628174650896 | 149,848416022847260 |
|  | 3 | ,213569323846160 | ,503209294670577 | . |
| *ACTA2* | 0 | ,657597178164632 | 10,480661509592496 | 487,450648788363300 |
|  | 1 | ,180203643401760 | 2,115735904334160 | 13,343710869520240 |
|  | 2 | ,294028988102860 | 3,577264215865210 | 38,216470564605310 |
|  | 3 | 1,306987923557221 | 12,223767759926512 | 36,692446501696820 |
| *IGF1* | 0 | 10,379487088528090 | 12,683615463628282 | . |
|  | 1 | ,143882081376972 | ,721724406684600 | 3,006978470722900 |
|  | 2 | ,028262648885863 | ,255788759061093 | ,753089264655963 |
|  | 3 | ,005447333342695 | ,039983109831595 | 2,183724712427208 |

| Gene | Macrophages | Q1 | Me | Q3 |
| --- | --- | --- | --- | --- |
| *MKI67* | 0 | ,227213446639651 | ,761765624399980 | 7,811207550629079 |
|  | 1 | ,399804528472078 | ,952853999385773 | 1,605666288256940 |
|  | 2 | ,232910086748650 | 6,478502926229186 | . |
| *ESR1* | 0 | ,203518348891032 | 1,405698435472364 | 4,534253923622811 |
|  | 1 | ,469356109489318 | 1,001844641963423 | 2,365807585711524 |
|  | 2 | ,817217416538734 | 3,611232569984251 | . |
| *PGR* | 0 | ,508794723202153 | 5,625346034188063 | 23,908477888141610 |
|  | 1 | ,527449370693254 | 16,219173607423090 | . |
|  | 2 | 26,616416507739128 | 52,195601452177730 | . |
| *TGFB1* | 0 | ,409697964259120 | 2,867334002157820 | 47,483594253543740 |
|  | 1 | 4,598429843148480 | 16,480628174650896 | . |
|  | 2 | ,213569323846160 | 75,030992673346700 | . |
| *ACTA2* | 0 | ,124573130093585 | 2,834666789353626 | 19,051549592482402 |
|  | 1 | 1,510717654855003 | 2,208299109137557 | 1240,576498047597600 |
|  | 2 | 1,878566748611178 | 24,984614527480100 | 178,032136048781700 |
| *IGF1* | 0 | ,037999954856177 | ,328894391521848 | 2,682777920427775 |
|  | 1 | ,003852833373386 | ,518287848896107 | . |
|  | 2 | ,206594453937480 | ,752034974942278 | 4,385960040226854 |

| Gene | Neutrophils | Q1 | Me | Q3 |
| --- | --- | --- | --- | --- |
| *MKI67* | 0 | ,141678804652306 | ,764255393930887 | 9,691832500989541 |
|  | 1 | ,473079332152158 | ,719434452011227 | 1,564272123347107 |
|  | 3 | 4,769347068044072 | 12,942722677889826 | . |
| *ESR1* | 0 | ,216182783438393 | 1,360351795757148 | 5,772436759063018 |
|  | 1 | ,439082638147788 | ,805446468124591 | 2,617659162451782 |
|  | 2 | ,000023320890304 | 1,190804860105994 | . |
|  | 3 | 3,777638559219151 | 4,155946241420981 | . |
| *PGR* | 0 | 1,511123647329750 | 6,897682244934275 | 26,616416507739128 |
|  | 1 | ,037347976860849 | 4,605533197781294 | 26,211133627480560 |
|  | 2 | ,527449370693254 | 82,047558172095240 | . |
|  | 3 | 1,030197266592703 | 7,840631800693917 | . |
| *TGFB1* | 0 | ,405505300459681 | 6,799706525829265 | 133,709249261409160 |
|  | 1 | 1,097394776270570 | 4,598429843148480 | . |
| *ACTA2* | 0 | ,379313104831280 | 3,577264215865210 | 32,815860233475930 |
|  | 1 | ,104298117721148 | 2,121776216906150 | 2,971859655090178 |
|  | 2 | ,002765093840898 | 2,104633559260825 | . |
|  | 3 | 1,728771736309773 | 68,621365922547060 | . |
| *IGF1* | 0 | ,121043025776710 | ,749926395514909 | 7,988026312020340 |
|  | 1 | ,001786697413002 | ,036599175308442 | . |
|  | 2 | ,003852833373386 | ,007352206898088 | . |
|  | 3 | ,007243547728959 | ,173203204633190 | . |

| Gene | Eosinophils | Q1 | Me | Q3 |
| --- | --- | --- | --- | --- |
| *MKI67* | 0 | ,241404927078880 | ,743089807736511 | 2,897476560929038 |
|  | 1 | ,072370139841806 | ,497337855574315 | 4,530928317037042 |
|  | 2 | 6,478502926229186 | 13,373063387214422 | . |
|  | 3 | 10,318707484443030 | 15,717402886089303 | . |
| *ESR1* | 0 | ,204367404449526 | 1,360351795757148 | 3,385263595837212 |
|  | 1 | ,250779864846500 | ,817217416538734 | 2,360547981174805 |
|  | 2 | 5,930218486733337 | 18,136527139336525 | . |
|  | 3 | 3,611232569984251 | 3,694435564601701 | . |
| *PGR* | 0 | ,527449370693254 | 5,541533368026351 | 25,284208184672604 |
|  | 1 | ,398768343876978 | 7,560227962215297 | 325,893229576514670 |
|  | 2 | 5,709158700349775 | 26,616416507739128 | . |
|  | 3 | 14,651066334795129 | 33,423333893486430 | . |
| *TGFB1* | 0 | ,581622167261982 | 5,078785920418415 | 44,733985133171060 |
|  | 3 | ,792849265494994 | 75,320632644171130 | . |
| *ACTA2* | 0 | ,315350017284965 | 3,150037332373271 | 21,018386899592270 |
|  | 1 | ,242308748484652 | 1,649905010284485 | 171,367419289342080 |
|  | 2 | ,002765093840898 | 7,391402358411920 | . |
|  | 3 | 1,728771736309773 | 2,653017976087492 | . |
| *IGF1* | 0 | ,082802771459447 | ,485077085431726 | 10,553936940480991 |
|  | 1 | ,019127527604728 | ,242251400992625 | 1,314189414316907 |
|  | 2 | ,069114639240400 | 2,832840087376495 | . |
|  | 3 | ,007243547728959 | ,380693551049303 | . |

| Gene | Irregular differentiation | Q1 | Me | Q3 |
| --- | --- | --- | --- | --- |
| *MKI67* | 0 | ,257701027182646 | ,764255393930887 | 10,701809941232323 |
|  | 1 | ,118358475386859 | 1,345043104775500 | 7,478031394529106 |
| *ESR1* | 0 | ,273338403110805 | 1,538337307847974 | 4,887417159360668 |
|  | 1 | ,204650422969024 | 1,315005156041931 | 3,611232569984251 |
| *PGR* | 0 | ,907115556748415 | 6,219607806480314 | 21,456505063708445 |
|  | 1 | ,404612349329392 | 27,512698734027698 | 34,916717190587360 |
| *TGFB1* | 0 | ,598478505677431 | 2,867334002157820 | 31,982111214097320 |
|  | 1 | ,409697964259120 | 36,485157772053020 | 1264,227808473060600 |
| *ACTA2* | 0 | ,120993361037474 | 2,538214723906008 | 27,282793573766526 |
|  | 1 | 1,052385821570234 | 5,484333287138565 | 18,363121343421970 |
| *IGF1* | 0 | ,028262648885863 | ,328894391521848 | 1,407902668777539 |
|  | 1 | ,048930379934637 | ,752034974942278 | 9,707106256132917 |

| Gene | Phase of the cycle 1-follicular 2-luteal | Q1 | Me | Q3 |
| --- | --- | --- | --- | --- |
| *MKI67* | 1 | ,127748457572242 | ,743089807736511 | 8,307410544638278 |
|  | 2 | ,563089721604770 | 1,564272123347107 | 7,478031394529106 |
| *ESR1* | 1 | ,239247504377131 | 1,007730116170494 | 3,652834067292976 |
|  | 2 | ,270877857857536 | 1,618552512618765 | 5,299091576052057 |
| *PGR* | 1 | ,530054634823828 | 5,176088978428975 | 22,785435769657003 |
|  | 2 | 1,164200795946685 | 11,333042758942707 | 32,526280441071230 |
| *TGFB1* | 1 | ,268019778702527 | 2,426727388230136 | 98,666005138195500 |
|  | 2 | 1,097394776270570 | 8,040271053970180 | 85,291748977094840 |
| *ACTA2* | 1 | ,169621921387987 | 2,107164575511497 | 17,497133967917980 |
|  | 2 | ,611819321435597 | 7,391402358411920 | 33,137566978020010 |
| *IGF1* | 1 | ,026038743045906 | ,518287848896107 | 11,531551276078186 |
|  | 2 | ,036599175308442 | ,442300508485868 | 1,710176269542400 |
